# Prokaryotic Diversity from Extreme Environments of Pakistan and its Potential Applications at Regional Levels

**DOI:** 10.1101/342949

**Authors:** Raees Khan, Muhammad Israr Khan, Amir Zeb, Nazish Roy, Muhammad Yasir, Imran Khan, Javed Iqbal Qazi, Shabir Ahmad, Riaz Ullah, Zuhaibuddin Bhutto

## Abstract

Extremophiles, the microorganisms thriving in extreme environments, provide valuable resources for practicing novel biotechnological processes. Pakistan homes a wide spectrum of extreme environments which harbor various biotechnologically significant microorganisms. This review gauges the structural and functional bacterial diversity of several extreme environments, emphasizing their potentials as a source of extremozymes, and in bioleaching, bioremediation, and bioenergy production at regional level. Further, this review highlights a panoramic account of the local natural conservatories of extremophiles. The inadequacies of current fragmental research are discussed with suggestions to quantitatively define the structural and functional diversity of unexplored extreme localities.

## 1. Introduction

Around half a century ago, people were largely unacquainted with the term extremophile, making it rather difficult to comprehend that life could exist at pH as low as 1, temperatures higher than 100 °C, high salinity, under 1400 atm atmospheric pressure and high concentrations of various recalcitrant substances [Hough and Danson, 199; Podar and Reysenbach, 2006]. However, like other discoveries, this hidden mystery was unmasked with the exploration of novel microbial communities – that remodeled the anthropocentric notions of habitable environments [MacElroy, 1974; Hendry, 2006]. Such microorganisms capable of both withstanding and surviving the extreme environments were distinguished thus named extremophiles [Podar and Reysenbach, 2006]. Subsequently, specific terms such as acidophiles, halophiles, thermophile, barophiles and psychrophiles representing the nature and corresponding habitat fascinated initially the microbiological and later biotechnological literature [Hendry, 2006; Cavicchioli et al., 2011]. An arsenal of specialized lipids, enzymes and proteins combat the austere constraints to fortify the structural and functional attributes of the cellular machinery in extremophiles [Berezovsky and Shakhnovich, 2005; Chen et al., 2005; Fukui et al., 2005; Falb et al., 2005; Choi et al., 2006].

The unique nature and interesting physiology of extremophiles piqued the interest of researchers and scientists to appraise the nature and dynamics of extremophiles and their derived components that could be a great frontier for biotechnological industry [Podar and Reysenbach, 2006]. The extremophile-derived components are already being applied in molecular biology, food industries, cosmetics, health sector, textile industries and waste processing units [Cavicchioli et al., 2011]. The need to study the abundance, activity, diversity and distribution of microorganisms in extreme environments at regional level for conserving their biodiversity and understanding the mechanisms of their survival in the unusual environments responding to the current global climate change are indispensable on cogent grounds. Furthermore, studies at regional level may strengthen our knowledge apropos of diverse nature of extremophiles inhabiting the same range(s) of extreme environment(s), but at different geographical locations.

Around the world, regional efforts have been carried out to evaluate structural and functional attributes of extremophiles’ diversity; for instance United States [Horikoshi, 1998], United Kingdom [Norton et al., 1993], India [Ghosh et al., 2003], China [Hu et al., 2015], Australia [Conner and Benison, 2013] and some African countries [Jones and Grant, 1999] are actively contributing to extremophile science. In this review, we focus on the extremophilic bacterial diversity and their habitats from various geographical regions of Pakistan discussing the so far explored extreme environments of the country and their reported bacterial diversity. Further, the review highlights possible biotechnological applications of locally isolated extremophiles for biotechnological industries. Moreover, new methods have been suggested for structural and functional evaluation of local virgin extreme sites.

### 1.2 Geographical importance of Pakistan

Pakistan is the 36th largest country and is located at 241° to 371° N latitudes and from 611° to 761° E longitudes. The total land area is about 8×10^5^ km^2^, which is expanded 1700 km from Northeast to Southwest with a total width of approximately 1000 km from East–West. The country has wonderful geomorphology that includes the lofty mountains of Himalayas, Karakorum, Hindukush, and Pamirs in the North with the fascinating coastline of the Arabian Sea in the South. In the central part of the country, mountains are bounded by the fertile plains of River Indus. One of the world’s largest deserts, known as Thar, is partially located in the Eastern part of the country. Also, the volcanic arc placed in Chagai, vast tectonic depression of Kharan. The westward swinging mountain ranges of Makran further enhance the importance of Pakistan [Zaigham et al., 2009]. Pakistan holds great geostrategic location with a diverse variety of natural resources that home many life forms. Various localities in the country, from which extremophiles were isolated, are summarized in Fig. 1 and Fig. 2.

**Fig. 1.**
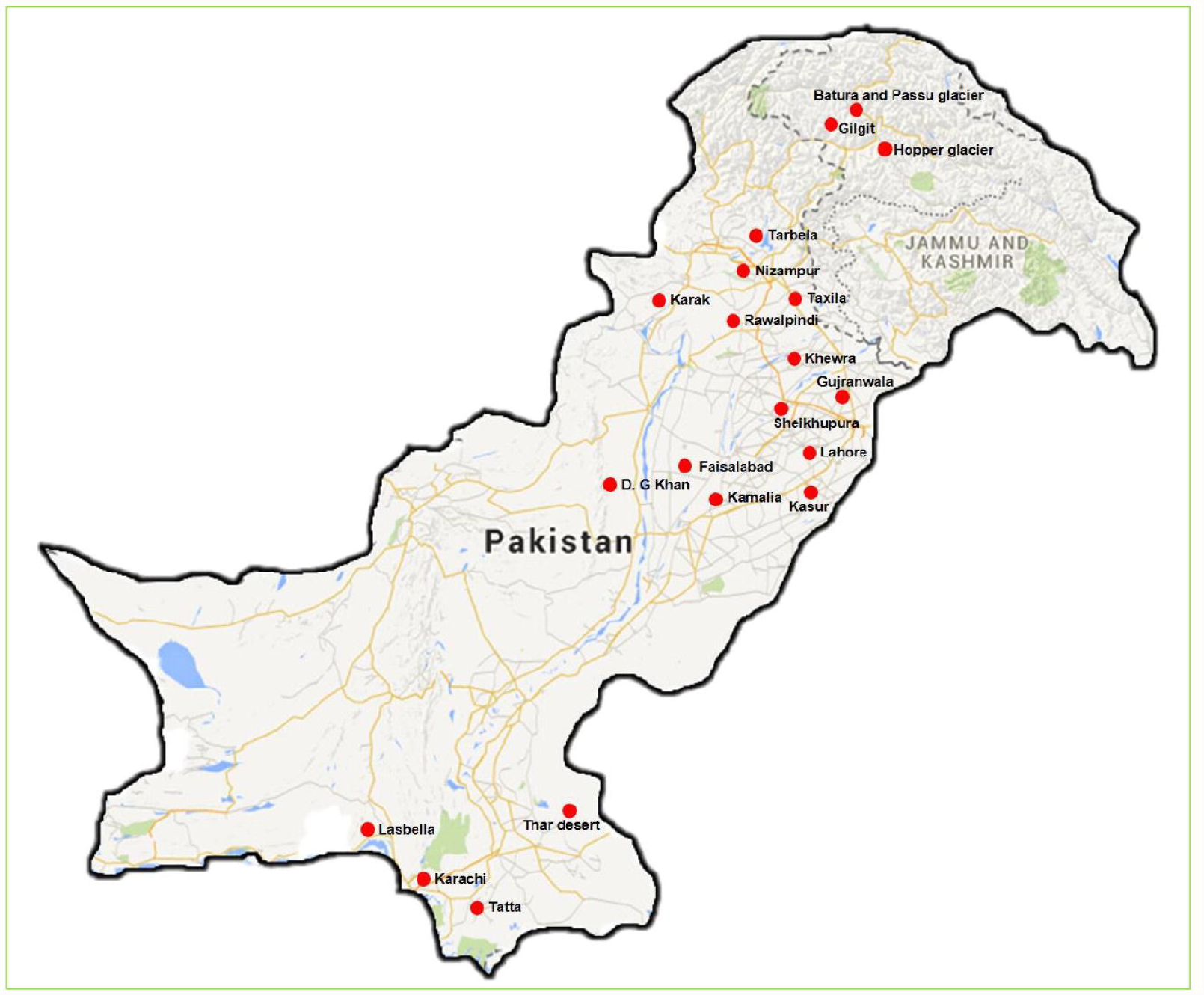
Map of Pakistan showing different localities from which extremophillic bacteria were isolated. The sample source area names are indicated with closed red circles.

**Fig. 2.**
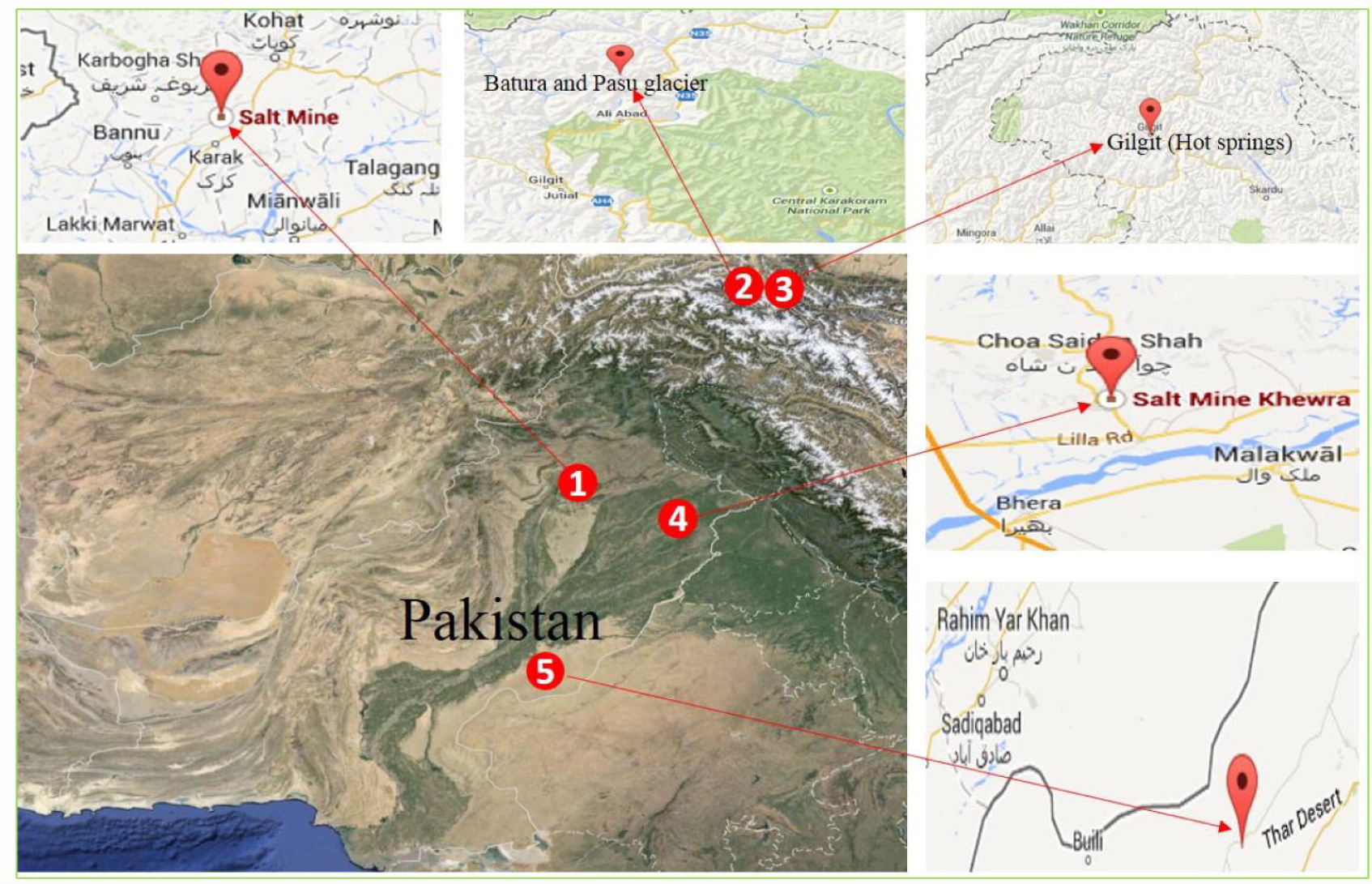
Map of Pakistan showing various natural extreme habitats of extremophiles in Pakistan, marked with closed numbered red circles. 1, 4; Salt mines of salt range, 2; various glaciers, 3; hot springs and 5; desert areas.

### 2.1 Thermophiles

Thermophiles are the organisms that dominantly inhabit hot environments [Urbieta et al., 2015]. These heat-loving microorganisms are homed in geothermally hot springs, hot soil, deep sea marine hydrothermal vents and sediments of volcanic islands [Mehta and Satyanarayana, 2013; Ebrahimpour and Kariminik, 2015]. They are also ubiquitous in hot polluted rivers and hot composts [Ebrahimpour and Kariminik, 2015]. Depending on the habitats thermophiles are classified into moderate (40–70 °C), extreme (above 70 °C) and hyper–thermophiles (80 – 105°C) [Ghosh et al, 2003; Reysenbach et al., 2002]. Thermophilic microorganisms are of immense importance in biotechnological research, as the organisms themselves and their derived products are widely used in food industry, textile industry, molecular research, bioleaching and pharmaceuticals [Ebrahimpour and Kariminik, 2015, Chaudhary and Qazi, 2008; Muhammad et al., 2009].

The global seismic belt passing through Pakistan results in the formation of volcanic and geo-pressurized thermal zone that sprouts out in the form of geothermal hot springs. Some of these hot springs are located in Murtazabad, Budelas, Tata Pani, Mashkin, Sassi and Chu Tran areas [Zaigham et al., 2009]. These hot springs are of local and national interest, as they are considered not only the potential source of renewable energy but also medicinally important [Javed et al., 2009]. The culture-based methods employed by few studies identified that *Bacillus*, *Staphylococcus*, *Streptococcus*, *Corynebacterium*, *Pseudomonas*, *Salmonella* and *Thermus aquaticus* species inhabit these springs [Javed et al., 2012; Saleem et al., 2012]. The thermophiles identified through culture-based methods from various environments of Pakistan are presented in Table 1. Interestingly, few industrially-beneficial strains such as *Thermus aquaticus*, *Geobacillus pallidus* sp. have been identified from these sites [Javed et al., 2012; Zahoor et al., 2012]. *Bacillus licheniformis* [Qadar et al., 2009; Niaz et al., 2010; Ghumro et al., 2011] and various other species of *Bacillus* have been also reported [Zaidi, 2007; Rafique et al., 2010; Khan et al., 2011; Asad et al., 2011]. These thermophilic isolates produce industrially important enzymes, including glucosidases, gelatin hydrolyses, extracellular α–amylases, cellulases, alkaline proteases, endoglucanases, and cellobiohydrolases. Majority of the enzymes have been reported stable at higher temperatures and salt concentrations. Additionally, some of the isolated thermophiles were capable of producing ethanol using sugarcane bagasse as a carbon source [Chaudhary and Qazi, 2008]. Few of the isolated thermophilic strains with antimicrobial properties could be further investigated as a potent source of therapeutic compounds [Muhammad et al., 2009]. Some of the isolated thermophiles were even capable to depyritize coal, oxidize iron and sulphur [Munawar et al., 2007]; whereas, others exhibited potential to leach out metals, (Table 1) [ Ilyas et al., 2007].

**Table 1.**
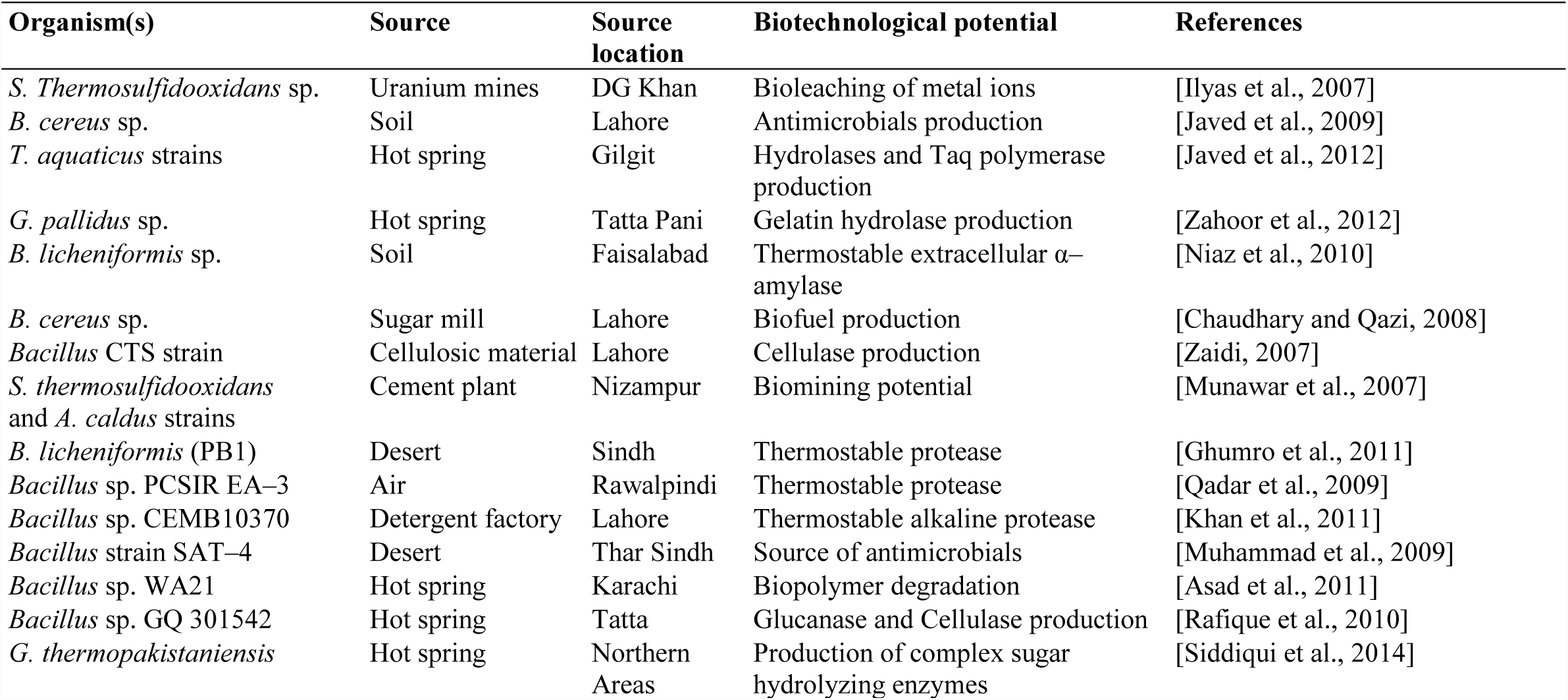
Thermophiles, isolated from various environments of Pakistan

### 2.2 Psychrophiles

Contrary to thermophiles, the psychrophiles grow and reproduce at low temperature ranging from 10 °C to −20 °C [Hough and Danson, 1999; Podar and Reysenbach, 2006]. Many psychrophilic species, belonging to both Gram–negative and Gram–positive bacteria, have been reported worldwide from various habitats including soil, fresh water marine lakes, sandstone, sea ice and oceans [Morita, 2000]. The psychrophilic bacterial enzymes are unique in catalytic activity and stability; having low activation energy that results in highly efficient catalytic reactions at lower temperatures [Podar and Reysenbach, 2006]. These bacteria may serve as excellent model systems to understand the molecular basis of low-temperature adaptation [Podar and Reysenbach, 2006].

Pakistan homes some of the world’s highest and most spectacular mountains. Surprisingly, 13 out of 30 world’s tallest peaks are located in Pakistan, including K2 (8,611 m), Nanga Parbat (8,125 m) and Tirich Mir (7,690 m) in the Hindu Kush mountainous range. Because of the numerous high mountains and abundant precipitation characteristic of a monsoon climate, the mountains of northern areas of Pakistan, including the Hindu Kush, Hindu Raj, Kohistan ranges, Nanga Parbat Massif and Karakoram Himalaya, has some of the largest and longest mid-latitude glaciers on earth. The glacierized area is estimated to cover 15,000 km^2^, and as much as 37 percent of the Karakoram region is covered by glaciers [Williams and Ferrigno, 2010]. The cold climate of this huge glacierized area might serve as a habitat for many cold-loving bacteria. Compared to the magnitude of diversity of the Psychrophylic environments the reports of bacterial diversity from such environments are scarce. However, a handful of the isolates reported from Batura, Hopper and Passu glaciers belong to genera of *Pseudomonas*, *Arthrobacter*, *Stenotrophomonas* and *Bacillus* [Ahmad, 2010] (Table 2). This calls for reconnaissance to help bring to light the structural and functional Psychrophylic diversity of remote and neglected environments of Pakistan. For this sake improved cultivation based techniques and metagenomics may be employed to peruse the phylogeny and functional capabilities of these organisms.

**Table 2.** Acidophiles and other extremophiles isolated from various environments of Pakistan

| Organism(s) | Source | Organism nature | Source location | Biotechnological potential | Reference |
| --- | --- | --- | --- | --- | --- |
| *L. bulgaricus*, *L. casei*, *L. acidophilus* and *L. salivarius* | Yogurt | Acidophile | Lahore | Probiotics and Dairy industry | [Aslam and Qazi, 2010] |
| *Bacillus* strain | Tannery waste | Acidophile |  | Lipase production | [Ghori et al., 2011] |
| *Bacillus* strain (RM16) | Hot spring | Acidophile | Karachi | Thermostable α-amylase | [Hassan et al., 2011] |
| *L. brevis*, *L. plantarum* and *L. fermentum* | Bakeries point | Acidophile | Faisalabad | Fermented bakery products | [Saeed et al., 2009] |
| *L. Paracasei* | Curd | Acidophile | Lahore | Dairy industry | [Ikramulhaq and Muktar, 2006] |
| *Acidithiobacillus ferrooxidans* | Black shale | Acidophile | Tarbela | Starch and gelatin hydrolysis | [Khan and Haq, 2012] |
| *T. thiooxidans*, *T. ferrooxidans* and *Azotobacter* | Water hyacinth | Acidophile | Taxila | Bioleaching of metals | [Mehmood et al., 2009] |
| *S. thermosulfidooxidans* | Coal heap | Acidophile | Faisalabad | Bioleaching of metals | [Ilyas et al., 2007] |
| *S. thermophiles* and *L. bulgaricus* | Homemade yogurt | Acidophile | Faisalabad | Mozzarella cheese production. | [Sameen et al., 2010] |
| *Methanosarcina mazei* and *Methanobacterium formicium* | Biogas plant | Methanophile | Karachi | Methane production | [Rajoka et al., 1999; Siddiqi and Robinson 1993] |
| Various *Bacillus* and *Klebsella* strains | Organic soil sample | Alkalophile | Kamalia | Leather processing and detergent industries | [Nadeem et al., 2007] |
| *Pseudomonas*, *Arthrobacter*, *Stenotrophomonas* and *Bacillus* generas | Glaciers | Psychrophile | Batura, Hopper and Passu glaciers | Alkaline phosphatase production | [Ahmad, 2010] |

### 2.3 Metallophiles

Metallophiles are the microbes that can withstand and tolerate environments characterized with high metal(s’) concentrations [Brakstad and Bonaunet, 2006; Head et al., 2006]. Mechanisms of metals’ uptake by these organisms, including biosorption and bioaccumulation processes, have got attention of the scientific community since long—because of their promising applications in the environment, agriculture, and industry [Tsezos, 2001]. A bracket of conceivable habitats of metalophiles may encompass natural mining sites, industrial biotopes, deep hydrothermal vents, volcanic areas and the nickel rich ultramafic soils [Mergeay, 2006]. Pakistan is gifted with various natural mineral reservoirs that include antimony, aragonite, barite, celestite, chromium, copper, gypsum, marble, salt and sodium compounds. Moreover, the largest iron deposits in the country are situated in western Punjab province with varying iron contents of 32–34% [Kuo, 2007]. Other ores include the low-grade uranium ores of Siwalik Baghalchur sandstone deposits in Suliman range [Khan, 1979]. Most of the reported metallophiles from Pakistan, however, have not been isolated from natural metallophilic environments. Therefore, it is necessary to harvest metallophiles from metallophiles habitats and to utilize their potentials. Metallophiles isolated from various environments of Pakistan are listed in Table 3. Metallophiles have adapted various approaches for coping with metals in corresponding biotopes. For instance, *C. freundii*, *K. oxytoca* and *B. anthracis* produce arsenic reductase enzymes that successfully reduce As(V) into As(III) and thus proceed to leach out arsenic [Shakoori et al., 2010]. Other metallophiles such as *S. thermosulfidooxidans* strains have been found to solubilize not only sulfide ores but other metals as well [Afrasayab et al., 2002]. Additionally a variety of metallophillic isolates such as *Pseudomonas* and *Sulfobacillus* spp., [Rehman et al., 2009], *Vibrionacea* sp. [Afrasayab et a., 2002], *T. thiooxidans*, *T. ferrooxidans* [Ilyas et al., 2007], *S. thermosulfidooxidans* and *A. ferrooxidans* [Rehman et al., 2009] were able to tolerate and leach out a variety of metals including uranium, chromium, arsenic, and mercury.

**Table 3.** Metallophiles isolated from various environments of Pakistan

| Organism(s) | Source | Source location | Biotechnological potential | References |
| --- | --- | --- | --- | --- |
| *Citrobacter freundii*,<br>*K. oxytoca* and *B. anthracis*<br>sp. | Wastewater plant | Sheikhupura | Arsenic reductase<br>production | [Shakoori et al., 2010] |
| *Pseudomonas lubricans* | Metal laden<br>industrial<br>wastewater | Sheikhupura | Heavy metal detoxification<br>potential | [Rehman et al., 2010] |
| *Sulfobacillus*<br>*Thermosulfidooxidans* | Uranium rich<br>environments and<br>salt mine | Khewra | Integrase enzymes<br>production and Uranium<br>leaching | [Ghauri et al., 2006;<br>Ghauri et al., 2003] |
| *Pseudomonas* sp. strains | Electroporating<br>units | Gujranwala | Chr (VI) detoxification | [Sultan and Hasnain,<br>2000] |
| *Vibrionacea* strains | Drainage<br>wastewater | Lahore | Environmental restoration<br>of Mercury | [Afrasayab et al., 2002] |
| *S. thermosulfidooxidans* and<br>*A. ferrooxidans* | Duddar deposit | Lasbela | Metal bioleaching,<br>biohydrometallurgical<br>potential | [Rehman et al., 2009] |
| *Bacillus* sp. KS21 | Industrially<br>contaminated sites | Kasur | Arsenic reduction | [Rehman et al., 2013] |
| *Cellulosimicrobium* sp. | Industrially<br>contaminated sites | Kasur | Chromium reduction | [Rehman et al., 2013] |
| *A. pakistanensis* | Textile dyeing<br>wastewater<br>treatment pond | Islamabad | Bioremediation | [Abbas et al., 2014] |

The biotechnological potentials of metallophiles from Pakistan can be utilized locally in a variety of ways; for instance, in commercial bioleaching operations and a wide range of bioremediation projects. Engineered bioleaching is already being used in many countries like Australia, South Africa, Chile, Myanmar, Ghana, Brazil and Peru for mining metals such as gold, cobalt and copper [ Brierley and Brierley, 2001]. Employment of endogenous metallophiles for bioleaching of low-grade ore heaps, such as those of iron and uranium, may pave the development of biohydrometallurgy in the country. Bioremediation of various metal contaminants, on the other hand, is of great interest amongst scientific communities [Xiong et al., 2008; Gadd, 2010; Guo et al., 2010; Monachese et al., 2012; Rajkumar et al., 2012; Basha and Rajaganesh, 2014]. A large number of industries in various cities of Punjab have led to the contamination of both ground as well as surface waters with pesticides, metals, and other toxic agents. The factories are dispensing toxic levels of Ca^2+^ Cl^−^, Ag^+^, Na^+^, K^+^, Mg^2+^, HCO^3-^, arsenic, mercury, iron, copper, lead, chromium, zinc, cadmium, cobalt and nickel [Azizullah et al., 2011]. Improper effluent disposals and misuses of agrochemicals have further exacerbated the problem. Merely a percent of these pollutants are properly treated prior to disposal [Azizullah et al., 2011]. There are various reports that these pollutants are badly affecting local people’s health [Azizullah et al., 2011; Arain et al., 2009; Farooqi, 2015]. Such cases call for an indispensable requisite to develop new, environment-friendly methods for effluent treatments to minimize environmental contamination. A potential approach to address the metal contamination is bioprocessing of the effluents. Mini scale biological waste water treatment plants, aided with metallophiles—if implemented in the local industries—may substantially reduce the metal load of wastewaters.

### 2.4 Halophiles

Halophiles are the extremophiles which flourish in saline habitats. This group of extremophiles is further categorized into mild, moderate and extreme halophiles on the basis of salt concentration of their habitats. The hyper-saline environments are diverse around the world and few of the well-known natural halophilic places include Lake Magadi (Kenyan Rift Valley), Owens Lake (California), Wadi Natrum Lake (Egypt) and several other saline soda lakes and soils [Oren, 2006; Qazi, 2013]. In addition to afore mentioned halophilic environments, coastal areas, deep-sea water, and underline salt mines are potential habitats for halophiles [Oren, 2006].

Pakistan is home to a variety of halophilic environments. Some of the local halophilic biotopes include the huge Salt range and mines of Khewra, Warcha and Kalabagh [Qazi, 2013], Noshpho [Khan et al., 2011], Uchhali Salt Lake [Hameed and Ashraf, 2008] and deep-sea environments Halophiles and alkalophiles, isolated from various halophilic environments of Pakistan are summarized in Table 4. From the Karak salt mine, different halotolerant strains belonging to various genera including *Oceanobacillus*, *Thalassobacillus, Terribacillus*, *Brevibacterium*, *Halomonas*, *Pseudomonas* and *Enterobacter* have been isolated utilizing culture based techniques [Roohi et al., 2012]. Few of the locally isolated halophilic strains have been analyzed to produce biotechnologically potent enzymes (Table 4). A halophilic *Bacillus* strain, isolated from Khewra salt range produced thermostable extracellular alkaline protease that was functional at 80 °C [Sehar and Hameed, 2011]. Other halophilic isolates from Khewra salt mine including *B. lichniformis*, *B. subtilis*, *K. oxytoca*, *B. megatarium*, *B. polymyxa*, *B. pumilis*, *B. brevis*, *B. macerans* and *B. coagulans* produced thermostable proteases which remained active over a broad range of alkaline pH [Akhtar et al., 2008]. Despite the interesting functional capabilities of these organisms and the diversity of halophilic extremophiles from Pakistani environments there still persists a dearth of reports about such isolates. To the best of our knowledge, not even a single report on the halophilic diversity of salt lakes and deep sea saline environment of the country has yet been reported which leaves a huge chasm for determining the structural and functional diversity of halophilic bacteria from these niches with robust culturomics and deep sequencing methodologies.

**Table 4.** Halophiles isolated from various environments of Pakistan

| Organism(s) | Source | Source location | Biotechnological potential | References |
| --- | --- | --- | --- | --- |
| *Bacillus pakistanensis* sp. | Salt mine | Karak | β-galactosidase, gelatin hydrolase and catalase production | [Roohi et al., 2014] |
| *B. fumarioli* | Salt mine | Khewra | Protease production | [Akhtar et al., 2008] |
| *B. sphaericus* | Salt mine | Khewra | Protease production | [Akhtar et al., 2008] |
| *B. amyloliquefaciens* | Salt mine | Khewra | Amylase, CMCase, Xylanase, Cellulase and Protease production | [Akhtar et al., 2008] |
| *B. cereus* | Salt mine | Khewra | Amylase, CMCase, Xylanase, Cellulase and Protease production | [Akhtar et al., 2008] |
| *S. arlettae* | Salt mine | Khewra | Amylase and Protease production | [Akhtar et al., 2008] |
| *S. gallinarum* | Salt mine | Khewra | Amylase production | [Akhtar et al., 2008] |
| *B.licheniformis* KL-176 | Salt mine | Khewra | Amylase, CMCase, Xylanase, Cellulase and Protease production | [Akhtar et al., 2008] |
| *B. pumilus* | Salt mine | Khewra | Amylase, CMCase, Xylanase, Cellulase and Protease production | [Akhtar et al., 2008] |
| 21 different halotolerant strains from various generas | Salt mine | Karak | Potential source of salt stress responsive genes | [Roohi et al., 2012] |
| *Cellulomonas pakistanensis* | Paddy grains | Islamabad | Plant growth promotion | [Ahmed et al., 2014] |
| *Kushneria pakistanensis* | Salt mine | Karak | Plant growth promotion | [Bangash et al., 2015] |

Abbreviations: CMCase; Carboxymethyl cellulase

### 2.5 Acidophiles

Acidophiles are the organisms that can grow and survive in acidic habitats [Canganella and Wiegel, 2011]. Such environments are particularly remarkable due to low pH (<5) causative of the microbial activity rather than the habitat itself [Gonzalez-Toril et al., 2013]. The Tinto River in southwestern Spain, a 100-km-long river; it has a low pH (between 1.5 and 3.1) and high concentrations of heavy metals in solution (iron at 0.4 to 20.2 g/L, copper at 0.02 to 0.70 g/L, and zinc at 0.02 to 0.56 g/L) is one such example of an extreme acidic ecosystem [Gonzalez-Toril et al., 2013]. The origin of acidic habitats has been considered to be associated with mining of metals and coal [Johnson, 1998]. The acidophiles usually thrive in natural or manmade metal rich habitats, mentioned afore in the metallophiles section. A variety of acidophilic bacterial isolates has been reported from several environments of Pakistan. Table 2 represents a summary of such micro-organisms. Among these acidophilic isolates, various *Lactobacillus* strains showed not only antagonistic activities against many pathogens but they may also be used in the production of good quality yogurt. Fermentation of animal feeds, acid tolerance, yogurt fermentation and antimicrobial characteristics of the bacterial isolates render them possible candidates for their use as probiotics [Aslam and Qazi, 2010]. Similarly, other acidophilic strains such as *L. brevis*, *L. plantarum*, *L. fermentum*, *L. paracasei* [Ikramulhaq and Muktar, 2006, Saeed et al., 2009], *S. thermophiles* and *L. bulgaricus* [Sameen et al., 2010] represent a potential source of starter cultures for different biological processes, such as production of fermented bakery products and good quality cheese with enhanced shelf life. Additionally, *S. thermophiles* and *L. bulgaricus* strains when compared with commercially available cultures for the production of mozzarella cheese showed faster acid production and efficient proteolysis which resulted in cheese with higher moisture content [Sameen et al., 2010].

The acidophilic extremophiles from Pakistani environments have a substantial economic potential to be utilized for production of fermented dairy products in the well-established dairy industry. Pakistan produces an estimated 42 billion liters of milk per annum [Shahid et al., 2012] and is the 5th largest producer of milk in the world. Though the industrial volume of dairy products could reach US$26 billion yet total milk production does not even suffice the domestic human needs, amounting to various problems [Shahid et al., 2012]. Utilizing the potentials of extremophilic microorganisms, especially those with probiotic characteristics and low or neutral pH stable enzymes in dairy fermentation processes, may improve the quality and availability of cost-effective fermented milk products, including cheese, yogurt and various other dairy products.

Some of the local acidophilic isolates also possessed bioleaching capabilities. For instance strains of *T. thiooxidans*, *T. ferrooxidans* and *Azotobacter* are capable of leaching out a variety of metals. Additionally, these strains have been found to play an important role in metals hyperaccumulation, nitrogen fixation and metal sulfide ores’ solubilization in their natural environments, such as in waste water and in soil (landfills) [Mehmood et al., 2009]. Another study showed that acidophilic heterotrophs such as *S. thermosulfidooxidans* were even capable of recovering metals from electronic scrap with leachabilities reaching up to 80% for Ni, Cu, Al, and Zn [Ilyas et al., 2007]. Moreover, coal biodepyritization studies conducted with pure and mixed consortia of some of the local acidophilic isolates revealed higher biodepyritization rates when such microbial consortia were employed [Munawar et al., 2007]. These local isolates may potentially be utilized in various bioleaching processes of low or high-grade ores. The potentials of these acidophilic extremophiles may also be utilized for recycling the local electronic wastes.

## 3. Potential applications of extremophiles at regional level-Recent trends

Since last decade, Pakistan has been facing a serious economic crisis due to war on terror, natural disasters, social and political instabilities and energy crisis. Taking energy crises into account, the current energy need of the country surpasses total generation [Khalil and Zaidi et al., 2014]. The country is mostly relying on hydel power, oil and nuclear power stations for electricity generation. These energy crises have rendered investors reluctant to invest and even driving the established industries to back out. The Pakistan Council of Renewable Energy Technologies (PCRE) was established to overcome the energy crises and to search and propose alternative energy sources; biofuel, for instance. The ministry of petroleum and natural resources of Pakistan, on the other hand, is the pioneer in the country that introduced biogas plants. From 1974 to 1987 the ministry installed 4137 biogas plants in different cities [Mirza et al, 2008; Bond and Templeton, 2011]. Later, in 2001 the PCRE technologies built 1200 more biogas plants [Bond and Templeton, 2011]. It is reported that a total of 5357 biogas plants are operational in the country [Bond and Templeton, 2011]. Most of these biogas plants harbor complex microbial communities, where the microbes of interest would have to compete for nutrients with other dominating groups of microorganisms which eventually limit the biogas production. Therefore, use of enriched pure cultures of biogas producing microorganisms would be of great interest in enhancing the yield of biogas plants. Despite the fact that the maintenance of pure cultures of these microbes would not be possible—as the substrate harbors a number of other species as well—still the higher number of the desired microbes will lead to comparatively efficient production of the biogas. This process would be further benefited if the continuous addition of pure cultures is maintained at various time intervals. To improve the productivity of the biogas plants, there is a need to shift from conventional methods towards recently used advanced approaches. In this instance, the huge number of already implemented biogas plants can be further profited with the use of extremophiles [Qurat-ul-Ain and Qazi, 2014]. Thermophilic processes have been correlated with better production of biogas [Angelidaki and Ellegaard L, 2003]. There is much likelihood that utilization of pure cultures of thermophilic microorganisms will enhance the efficiency of the biogas plants. Some of the local extremophiles have been reported capable of producing methane gas (Table 2), they can potentially be employed in biogas plants. The sugar mills industry of Pakistan stands as another sector, where utilization of such processes has great potentials for methane gas production through bagasse degradation. These sugar mills utilize 4670000 tons of sugarcane, leaving an average 1401000 tons of bagasse [Rehman MS, 2013], which can potentially be used to generate 2000 MW of electric power [Mirza et al., 2008]. Meanwhile, only two out of 78 sugar mills of the country, namely the Shakar Ganj Sugar Mills (Jhang) and Habib Sugar Mills (Nawabshah), have adopted such processes for the production of biofuel. Both the mills generate sufficient electricity by growing biogas producing microbes on bagasse, that fulfills not only the energy needs of the sugar mills but even they sell the surplus electricity to the national grid [Malik, 2014]. It is being expected from both the government and the private sector to utilize biotechnological approaches to achieve the country’s energy demands. The Pakistan sugar mills association should convince the fellow industrialists to implant such technologies that are cost-effective and environment-friendly.

Beside biofuel applications, microorganisms synthesize valuable enzymes that are routinely used in industries like textile, leather, hosiery, and denim. The enzymes of extremophiles are resistant to the extreme processing conditions of industry. To meet the industrial demands of the country, around 500 tons of cellulase and 5000-7000 tons of α-amylase are imported each year costing over $10 million every year on the import of dextransucrase, α-amylase, and protease. These enzymes could be locally produced which in turn can reduce the expenses of industry and generate more jobs [Dawn News, 2009]. This can be achieved by tapping the potential of producing cheap agriculture commodities such as biotechnologically potent enzymes from local bacterial isolates utilizing agricultural wastes (tables 1– 4). Keeping the industrial demand for enzymes, the country is investing in enzyme research sector. The trend for a search of beneficial microorganisms capable of producing biotechnologically potent enzymes has been on the rise. Various, enzyme-research oriented projects are currently running in most of the biotechnological institutes of the country. Recently, an enzyme has been isolated from the hyperthermophilic archaeon *Thermococcus kodakaraensis* KOD1 with promiscuous pullulanase and α-amylase activities. Additionally, the enzyme was stable at higher temperatures (95-100°C) and a broad range of pH (3.0-8.5). The enzyme has been patented (US 20140227744 A1) and the researchers are interested in its bulk production. National Institute of Biotechnology and Genetic Engineering (NIBGE), being one of the leading biotechnological research institutes of the country, is currently focusing on extremophiles as a source of novel and biotechnologically potent enzymes [Ghaffar et al., 2011]. Researchers from industrial biotechnology group at NIBGE have isolated several other enzymes that have promising applications in pulp, bio-stone washing and bio-finishing of garments. Several fiber-degrading enzymes are in the process of commercialization to be applied in the poultry and ruminant feed industries (personal communications and NIBGE website). Additionally, NIBGE has taken steps forward for the qualitative and quantitative improvement of enzymes, utilizing various approaches, such as mutagenesis [Arshad et al., 2014] and solid state fermentation [Tabassum et al., 2014]. Similarly, Dr. Abdul Qadeer Khan Institute of Biotechnology and Genetic Engineering (KIBGE) of Karachi University recently announced the mass scale production and commercialization of five industrially important enzymes, including dextransucrase, α-amylase and protease [Dawn News, 2009]. Several other universities and research centers of the country are also trying to optimize conditions for mass level production of various enzymes. However, NIBGE and KIBGE are currently the leading institutes in enzyme research in Pakistan.

## 4. Conclusions and perspectives

Pakistani extreme environments, being diverse, unique and rich in terms of abiotic and biotic components home a variety of extremophiles. The conventional based approaches utilized so far to study the microbial diversity of these extreme environments are not sufficient lacking even a single report on the structural based diversity of any of the extreme environments of the country. Moreover, some of the extremophilic isolates from these environments are not true extremophiles because extremotolerants are usually confused with extremophiles by the local graduates. Additionally, some of the extreme environments of the country have been completely disregarded so far (e.g. Peatlands). There are numerous peatlands in northern areas of Pakistan, such as Deosai peatland, located in Skardu, Gilgit-Baltistan (Latitude: 35° 03′ 40″ N and Longitude: 75° 28′ 34″ E). This peatland is situated at an average elevation of 13,497 feet (4,114 M) above the sea level, which makes it one of the highest plateaus in the world. Other unexplored extreme environments include the enormous number of caves in Pakistan. To date, there are more than 100 recorded caves in Pakistan [Gunn, 2004]. Few examples of caves from Pakistan include “Pir Ghaib Gharr Gharra” Balochistan (1,275 M), “Murghaghull Gharra” Balochistan (576 M) and Kach Gharra (Juniper shaft Cave) Balochistan (131 M). Microbial diversity is considered as a function of the environment, therefore these unexplored environments might harbor biotechnologically potent microbes and unique diversity. Exploration of extremophiles with modified culture conditions and structural or functional based metagenomic approaches may lead to the isolation of novel microbes and may explore their biotechnological potentials [Vester et al., 2015]. The above-narrated strategies will help to isolate new extremophiles that may encode industrially valuable enzymes. Most of the microorganisms in nature are not culturable, and one might imagine the diversity and presence of novel genes for various biotechnologically potent enzymes from the majority of unculturable microorganisms. Function-driven metagenomics is an effective way to search for novel genes and enzymes from the diverse microorganisms. Such biotechnological approaches may help to domesticize extremophiles from their inhabitable natural locations and the efforts may bring oversized impact in our lives.

## 6. Author Contributions

RK, MK and AZ. Through the idea and conceived and organized the manuscript. RK, MK, AZ, IK, YC, NR, SA, RU and ZB interpreted and analyzed the data and prepared the manuscript. RK, MY, and JQ, critically evaluated the manuscript. All of the authors read and approved the final version of the manuscript before submission.

## 8. List of abbreviations

PCRE: Pakistan Council of Renewable Energy Technologies
NIBGE: National Institute of Biotechnology and Genetic Engineering
KIBGE: Dr. Abdul Qadeer Khan Institute of Biotechnology and Genetic Engineering

## Acknowledgments

We thank Dr. Junaid Khaliq for critically reading this paper.

